# Cabozantinib unlocks efficient *in vivo* targeted delivery of neutrophil-loaded nanoparticles into murine prostate tumors

**DOI:** 10.1101/2020.04.13.037531

**Authors:** Kiranj Chaudagar, Natalie Landon-Brace, Aniruddh Solanki, Hanna M. Hieromnimon, Emma Hegermiller, Wen Li, Yue Shao, Devan Wilkins, Kaela Bynoe, Xiang-Ling Li, John Clohessy, Soumya Ullas, Jeffrey M Karp, Akash Patnaik

**Author notes:** N.L.B., K.C. and A.S. have contributed equally to this manuscript as first authors. J.M.K and A.P. have contributed equally to this manuscript as senior authors. Corresponding Author: Dr. Akash Patnaik, MD, PhD, MMSc, The University of Chicago, Knapp Center for Biomedical Discovery, Room 7152, 900 East 57^th^ Street, Chicago, IL 60637.

## Abstract

A major barrier to the successful application of nanotechnology for cancer treatment is the efficient delivery of therapeutic payloads to metastatic tumor deposits. We have previously discovered that cabozantinib, a tyrosine kinase inhibitor, triggers neutrophil-mediated anti-cancer innate immunity, resulting in tumor regression in an aggressive PTEN/p53-deficient genetically engineered murine model of advanced prostate cancer. Here, we specifically investigated the potential of cabozantinib-induced neutrophil activation and recruitment to enhance delivery of bovine serum albumin (BSA)-coated polymeric nanoparticles (NPs) into murine PTEN/p53-deficient prostate tumors. Based on the observation that BSA-coating of NPs enhanced association and internalization by activated neutrophils *in vitro*, relative to uncoated NPs, we systemically injected BSA-coated, dye-loaded NPs into prostate-specific PTEN/p53-deficient mice that were pre-treated with cabozantinib. Flow cytometric analysis revealed a 4-fold increase of neutrophil-associated NPs within the tumor microenvironment (TME) of mice pre-treated with cabozantinib relative to untreated controls. At steady-state, following 3 days of cabozantinib/NP administration, 1% of systemically injected dye-loaded NPs selectively accumulated within the TME of mice that were pre-treated with cabozantinib, compared to 0.11% uptake for mice that did not receive cabozantinib pre-treatment. Strikingly, neutrophil depletion with Ly6G antibody abolished NP accumulation in tumors to baseline levels, demonstrating targeted neutrophil-mediated NP delivery to the prostate TME. In summary, we have discovered a novel nano-immunotherapeutic strategy for enhanced intratumoral delivery of injected NPs, which results in significantly higher NP accumulation than reported strategies in the nanotechnology literature to-date.

## Introduction

Prostate cancer (PCa) is the most commonly diagnosed cancer in the United States, with an estimated 160,000 new diagnoses each year (1). An estimated 1 in 6 men will be diagnosed with PCa in their lifetime, most with localized disease. Metastatic PCa accounts for 29,430 deaths in the US annually, with 5-year survival rates at less than 29% (1,2). The majority of men with metastatic PCa will develop resistance to primary androgen deprivation therapy, leading to metastatic, castration-resistant prostate cancer (mCRPC) (3). Several therapies have been FDA approved for mCRPC over the last 10 years. These include sipuleucel-T vaccine, androgen-receptor targeted agents like abiraterone and enzalutamide, taxane chemotherapy and Radium-223, that improve overall survival in mCRPC (4). However, most patients still develop therapeutic resistance, highlighting an unmet need to establish more potent, curative treatments with minimal toxicity (5).

In recent years, there has been considerable success of immune checkpoint blockade (ICB) therapy across multiple cancers (6). However, there are still large subsets of patients across multiple cancers that do not respond to these therapies. The responses of mCRPC to ICB have been limited, highlighting the critical need to design novel therapeutic strategies that can harness the immune system to enhance therapeutic efficacy (7). We have previously demonstrated that cabozantinib, a promiscuous receptor tyrosine kinase (RTK) inhibitor, FDA approved in medullary thyroid, renal, and hepatocellular carcinoma, unleashes a potent neutrophil-mediated innate immune response, resulting in tumor eradication in mice with probasin-Cre driven conditional prostate-specific knockout of PTEN/p53 (Pb-Cre; PTEN^fl/fl^p53^fl/fl^) (8). This study demonstrates the potential of cabozantinib to reprogram neutrophil-mediated innate immunity to eradicate advanced CRPC, and has led to >20 combination clinical trials of cabozantinib and ICB across multiple malignancies. In particular, a recent Phase II clinical trial evaluating cabozantinib and atezolizumab (COSMIC 021) in multiple different tumor types, demonstrated an overall response rate of 32%, disease control rate of 80% and a PSA decline rate of 50% in the cohort of men with CRPC (9). Based on these promising results, a Phase III registration trial of this combination is currently being planned in mCRPC.

Over the past decade, nanoparticles (NPs) have been investigated as a drug delivery system to enhance chemotherapeutic concentrations within tumors, while minimizing off-target toxicity (10). However, efficient NP delivery has been a major challenge for clinical translation, with studies indicating that 0.7% (median) of injected NPs are actually delivered to the tumor (11). Polymeric NPs made of materials such as poly(lactic-co-glycolic acid) (PLGA) are a popular choice for chemotherapeutic delivery due to their tunability, versatility and ability to provide controlled drug release (12). NPs often rely on the enhanced permeability and retention effect in tumors for delivery, resulting in NPs becoming trapped away from their intended target and preventing efficient delivery of chemotherapeutic agents to tumor cells (13). While several approaches have been explored to improve NP delivery using active cellular targeting, the majority have not demonstrated success in clinical trials (14). Therefore, a critical need exists to improve the targeting and delivery of polymeric NPs to tumor deposits.

In this study, we tested the hypothesis that cabozantinib-mediated neutrophil activation/infiltration will result in enhanced delivery of systemically injected NP into the prostate tumor bed. Prior studies have used bovine serum albumin (BSA) to enhance internalization of NPs into neutrophils, which can then carry NPs to the tumor bed (15-17). In this study, we utilized PLGA-based NPs coated with native BSA (see below), which offer several advantages over BSA-only NPs, including their ability to encapsulate a wider variety of agents, the ease of tuning size and drug-loading, and the ability to modify drug release rates based on application. Here we designed PLGA NPs coated with native BSA (commonly referred to as BSA) to achieve neutrophil-specific delivery to prostate tumors. BSA-coated, dye-loaded PLGA NPs (BSA-NPs) were injected systemically into genetically engineered mice that develop prostate tumors as a result of prostate-specific PTEN and p53 loss (Pb-Cre; PTEN^fl/fl^p53^fl/fl^ mice). We observed that accumulation of dye-loaded NPs in the tumor was enhanced in mice pre-treated with cabozantinib, relative to untreated controls. Furthermore, the enhanced intratumoral NP accumulation with cabozantinib pre-treatment was abrogated via concomitant administration of Ly6G antibody, which depletes neutrophils, suggesting that NP delivery is neutrophil-mediated. This novel nano-immunotherapeutic strategy has the potential to deliver cancer medicines with narrow therapeutic indices (low efficacy/toxicity ratios), thus overcoming current challenges to passive and active targeting of NPs into tumors.

## Materials and Methods

### Preparation & Characterization of Nanoparticles

PLGA NPs were prepared by single emulsion as previously described (12). 50 mg of 50:50 poly(DL-lactic-co-glycolic)-COOH (Lactel Absorbable Polymers, cat. B6013-1) and 10 μL of 1 mM Vybrant(R) Cell-Labeling DiO or DiR dye (ThermoFisher, cat. V22889) was added to 2 mL of dichloromethane (Sigma Aldrich). Once PLGA/dye was fully dissolved, the solution was sonicated for 10 seconds before being added dropwise to 1% wt/vol solution of filtered poly-vinyl alcohol (Sigma Aldrich), while homogenizing at 35000 rpm for 2 minutes. Particle suspension was stirred and organic solvent was allowed to evaporate for 4 hours. Particle suspension was washed 3 times with 10 mL of distilled water by centrifuging solution in 15 mL centrifuge tubes at 1000g for 5 minutes. After the final wash, particles were resuspended in 1 mL of distilled water and characterized by dynamic light scattering for size and zeta-potential. For coating, NPs were incubated at 5 mg/mL in a 20 μg/mL Bovine Serum Albumin (BSA; Sigma-Aldrich) solution for 2 hours at 37°C. NPs were washed twice with distilled water at 1000g for 5 mins and resuspended in PBS at the desired concentration. BSA coating was verified using UV spectroscopy.

### *In vitro* Studies

For human experiments, near-confluent PC3 cells cultured in RPMI-1640 with 0.1% BSA and 1% penicillin/streptomycin, were treated with 5 uM cabozantinib for 24 hours. The supernatant was collected and utilized as PC3-conditioned media, to activate human neutrophils for *in vitro* experiments. For murine experiments, near-confluent PTEN/p53-deficient prostate tumor-derived SC1 cells cultured in PrEGM Bulletkit media supplemented with 10% FBS and 1% penicillin/streptomycin, were treated with 10 uM cabozantinib for 24 hours. The supernatant was collected and utilized as SC1-conditioned media to activate and murine neutrophils.

### Isolation of Neutrophils

Human neutrophils were isolated from 10 mL of human whole blood obtained from Research Blood Components (Boston, MA). Informed consent was obtained from blood donors by Research Blood Components prior to collection. The EasySep™ Direct Human Neutrophil Isolation Kit was used as per standard protocol (StemCell Technologies, cat. 19666). Isolation cocktail containing antibodies and RapidSpheres™ were added to the whole blood sample and immunomagnetic negative selection was performed. Neutrophils were collected in serum-free RPMI-1640 (Gibco) with 1% penicillin/streptomycin for *in vitro* studies. Neutrophil population was verified using anti-CD11b antibody (Biolegend 101257) and anti-CD16 antibody (Biolegend 302025) staining by flow cytometry.

Murine neutrophils were isolated from 500 uL of whole blood obtained from Pb-Cre; PTEN^fl/fl^, p53^fl/fl^ mice, in accordance with NIH guidelines and protocol approved by the IACUC at University of Chicago. The EasySep™ Mouse Neutrophil Enrichment Kit was used as per the included protocol (StemCell Technologies, cat. 19762). Isolation cocktail containing antibodies and RapidSpheres™ were added to the whole murine blood sample and immunomagnetic negative selection was performed. Neutrophils were collected in phosphate buffer saline pH 7.4 (Corning) with 2% fetal bovine serum (FBS, Gemini) and 1mM ethylenediaminetetraacetic acid (EDTA, Thermofisher Scientific) penicillin/streptomycin for *in vitro* studies. Neutrophil population was verified using anti-CD11b antibody (Biolegend 101257) and anti-GR1 antibody (Biolegend 108422) staining by flow cytometry.

### *In Vitro* Human Neutrophil Activation/NP Internalization/Imaging

Neutrophils were activated by incubating with PC3-conditioned media on an 8-chamber slide at 37°C for 3 hours. Following this incubation, the non-adherent neutrophils were washed twice with PBS. 100μg of BSA-coated DiO-loaded PLGA NPs were suspended in PC3-conditioned media and incubated with adherent neutrophils on an 8-chamber slide at 37°C for 1 hour. Media was removed and chambers were washed once with PBS. Neutrophils were stained with DiI (1:1000, ThermoFisher Scientific) at 37°C for 15 minutes to delineate the plasma membrane. Neutrophils were washed three times with PBS at 37°C for 10 minutes to remove excessive DiI dye. Next, nuclei were stained with Draq5 (1:1000, ThermoFisher Scientific) at 37°C for 5 minutes and imaging was carried out using a Leica SP5-STED microscope. The internalized NPs appeared green and surface-bound NP appeared fluorescent red-yellow in color.

### *In Vitro* Murine Neutrophil Activation/ImageStream

Neutrophils were activated by incubating with SC1-conditioned media at room temperature for 15, 60, 120 and 180 minutes. For the aforementioned times, neutrophils were incubated with 10, 100 and 1000μg of BSA-coated DiR-loaded PLGA NPs in 1.4 mL of conditioned media at room temperature. Neutrophils were stained with DAF-FM diacetate (ThermoFisher Scientific D23844) at room temperature for 15 minutes before completion of incubation period to assess nitric oxide (NO) level per neutrophil. Following incubation, neutrophils were analyzed for associated NP and NO using ImageStream Mark II (Amnis; Luminex). Neutrophils were passively bound to 1 or 2 BSA-NP, whereas activated ones could bind >2 BSA-NP. MFI of NP were assessed only for activated (or more than 2NP associated) neutrophils.

### *In vitro* Murine Neutrophil Cytokine assay

Neutrophils were activated by incubating with 1.4 mL of SC1-conditioned media containing 0, 10, 100, 1000μg of BSA-coated DiR-loaded PLGA NPs at room temperature for 6 hours. After incubation period, neutrophils were centrifuged and supernatants were collected for estimation of neutrophil-secreted cytokines. Cytokine array kits containing anti-viral response panel (Cat no. 740622) and proinflammatory chemokine panel (Cat no. 740451) were used according to manufacturer protocol (LegendplexTM platform, Biolegend).

### *In vitro* Murine Neutrophil Phagocytosis assay

SC1 cells were treated *in vitro* with cabozantinib for 24 hours, and media was replaced with DiI dye (ThermoFisher, Cat. no. V22885) containing PBS for 20 minutes, to stain membranes of tumor cells. Following aspiration of dye, SC1 cells were next washed with PBS (3x) to completely remove dye within wells. 500,000 neutrophils were suspended in SC1-conditioned media containing 0, 10, 100, 1000μg of BSA-coated DiR-loaded PLGA NP and further added on the top of DiI dye-stained SC1 cells. After 6 hours incubation, anti-CD11b antibody (Biolegend 101257) and anti-GR1 antibody (Biolegend 108422) were added at dilution of 1:100 in wells for 30 minutes. Neutrophils were further fixed and lifted by trypsinization to determine DiI dye per neutrophil using flow cytometry (18,19). Percent phagocytosis was calculated by scaling MFI of DiI per neutrophil in presence and absence of SC1-conditioned media to 100% and 0%, respectively.

### *In Vivo* Studies

Pb-Cre;PTEN^fl/fl^p53^fl/fl^ mice were screened for prostate tumor development at 16 weeks of age by ultrasound. Following the development of solid tumors, the mice were treated with cabozantinib (100 mg/kg/day, oral gavage) when the tumors reached a long-axis diameter of 10 mm. One hour after cabozantinib treatment, 50 μg of BSA-coated or uncoated DiO-loaded PLGA NPs and 50 μg of BSA-coated or uncoated DiR-loaded PLGA NPs (total 100 μg NPs) were intravenously injected every 12 hours for 3 days. Mice were sacrificed at 72 hours and peripheral blood and the following tissues were collected: prostate, liver, kidney, spleen, lungs, and femur/tibia. Imaging was conducted using the *In Vivo* Imaging System (IVIS) and processed by spectral unmixing to control for background signal and capture fluorescence from DiR particles. These fluorescent images were then overlaid with bright field images of the organs for qualitative and quantitative analysis. Images were analyzed with ImageJ, and mean fluorescence was obtained for each tumor for comparison.

### Flow Cytometry

For neutrophil/NP association analysis within tumor and spleen, tissues were homogenized using liberase, and filtered with HBSS through a 70-μm cell strainer. The resulting cell suspension was centrifuged and incubated for 2 mins in 1 mL of ACK solution to lyse red blood cells. ACK was neutralized with 10 mL of HBSS and the cell suspension was centrifuged at 1800 rpm for 5 minutes. This was repeated 3 times or until no red cells were visible (whichever came first). The final cell pellet was resuspended in 5 mL of HBSS and distributed in equal volumes for flow cytometry. Cells were centrifuged at 1800 rpm for 5 mins and then incubated in 2 μg/mL solution of anti-CD11b antibody (Biolegend 101257) and anti-GR1 antibody (Biolegend 108422) for 30 minutes in the dark. Cells were washed once with HBSS and run on BD Fortessa 4-15 flow cytometer until 10,000 events were captured in the population gated for mouse neutrophils. For analysis of peripheral neutrophil/NP association, blood was incubated for 2 minutes in 1 mL of ACK solution to lyse red blood cells, then centrifuged and the supernatant was aspirated. The pellet was subsequently stained with anti-CD11b antibody (Biolegend 101257) and anti-GR1 antibody (Biolegend 108422) for flow cytometry analysis of NP-associated circulating neutrophils.

### Nanoparticle uptake determination

A 50 mg sample of harvested tumors was homogenized in 1 mL PBS then pelleted at 35,000 rpm for 1 min, and this step was repeated a total of three times. The concentrations of DiO NPs were measured in homogenates at 485 nm excitation and 510 nm emission wavelengths using a fluorometer. Standard DiO NP solution was prepared using tumor homogenates of mice that were not treated with NPs and cabozantinib. The linearity range was 2-20 ng DiO NPs/uL (R=0.9674). The % NP uptake was calculated based on a single administered dose of NPs.

### Data Analysis

Unless otherwise indicated, data was analyzed using GraphPad Prism 7 (GraphPad Software Inc.) and statistical analysis was performed using unpaired student t-test with p < 0.05 level of significance.

## Results

### BSA-coating of NPs enhanced association and internalization by activated neutrophils *in vitro*

Figure 1A shows a schema for the design of BSA-NPs utilized in this study. DiO- and DiR-loaded NPs were initially characterized by Dynamic Light Scattering (DLS). DLS revealed an average particle size of 450 nm with average size ranging from 400-500 nm across batches. These particles are significantly larger than the 100-200 nm BSA-coated PLGA NP used in prior studies (20), and may allow for increased drug loading with sustained release of drug(s) per unit NP (15,16). The NP surface charge, pre- and post-BSA coating, was on the order of −40 mV, consistent with previous studies of the expected charge of ovalbumin containing PLGA NPs. Scanning electron microscopy (SEM) confirmed the size distribution and surface morphology of NPs, which was on the order of 500 nm, consistent with DLS measurements. Furthermore, SEM showed smooth, spherical particles with minimal aggregation (Figure 1B).

**Figure 1.**
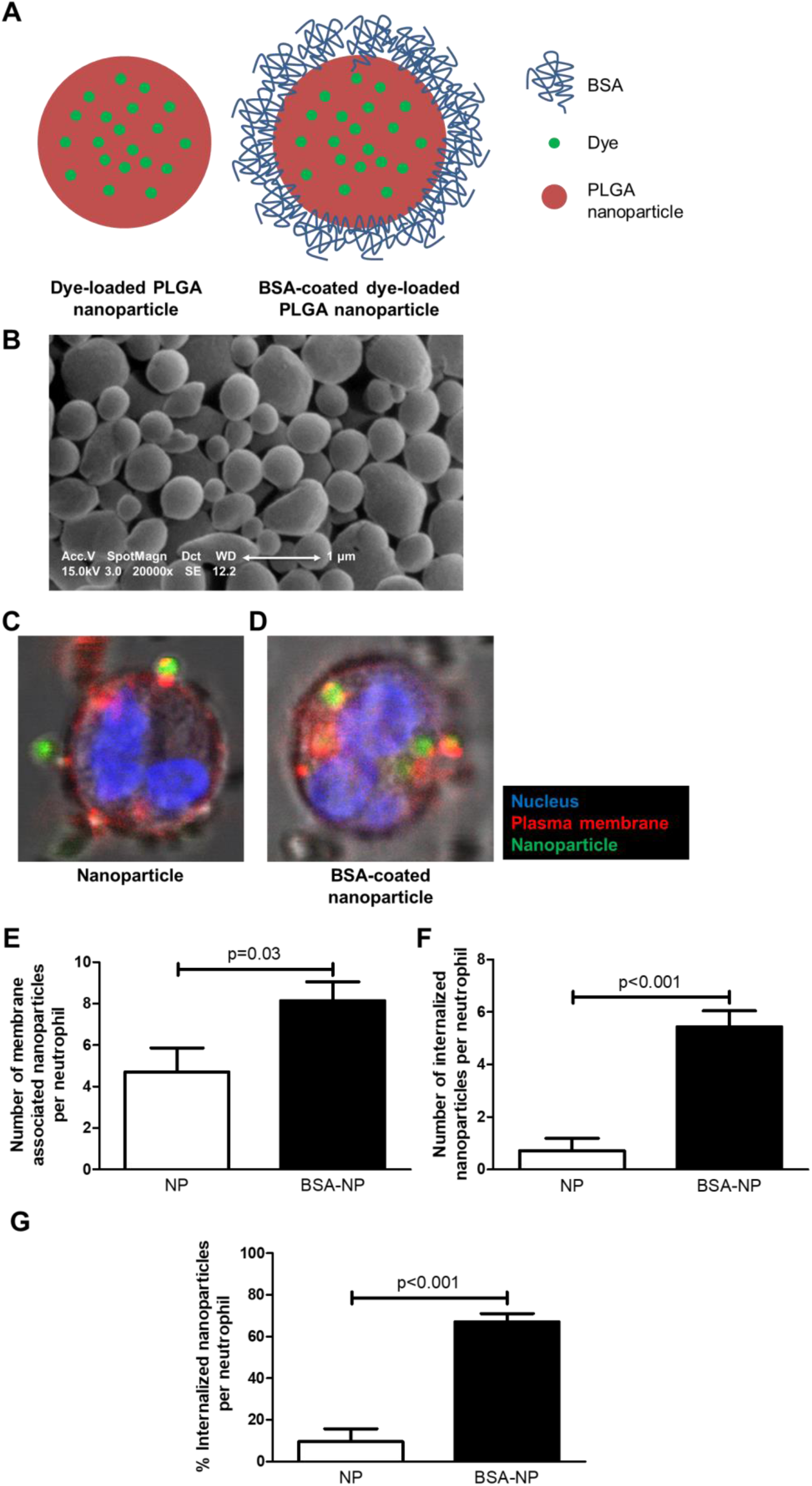
BSA-coated PLGA NPs (BSA-NPs) preferentially attach to activated human neutrophils *in vitro*, relative to uncoated NPs. (A) Schema illustrates proposed dye-loaded PLGA nanoparticle with and without BSA coating. (B) Scanning electron microscopy (SEM) performed to assess NP size and shape. (C, D) *In vitro* internalization of BSA-NPs by neutrophils. Human PC3 prostate cancer cells were treated with cabozantinib, and conditioned media was utilized to activate neutrophils followed by incubation with BSA-coated or uncoated PLGA NPs. Cells were gently washed with PBS before imaging under confocal microscope to remove unassociated NPs. (E-G) Quantification of the average number of NPs associated with neutrophils for BSA coated and uncoated NPs was performed by counting the number of visible DiO particles and dividing by the number of nuclei in representative images per condition. The internalized NPs appeared green and surface-bound NP appeared fluorescent red-yellow in color. The % internalized NPs per neutrophil was calculated by the ratio of membrane associated (red-yellow fluorescent)/internalized (green) nanoparticles X 100. n=3 independent experiments.

To determine whether BSA-coated NPs would preferentially associate with and be internalized by activated neutrophils relative to uncoated NPs, we performed an *in vitro* neutrophil activation assay. Neutrophils were activated *in vitro* with conditioned media from human PTEN/p53-deficient PC3 cells, and incubated with BSA-coated, DiO-loaded PLGA NPs. Using confocal microscopy, an increased number of DiO-loaded NPs were found to be associated with, and internalized by, activated neutrophils when pre-coated with BSA. An average of 8.15 ± 1.14 BSA-NPs were associated with activated neutrophils, compared to an average of 5.3 ± 0.7 uncoated NPs associated with activated neutrophils (p < 0.05). BSA coating resulted in an approx. 6-fold enhancement of the internalization of NPs (5.46 ± 0.92 internalized NPs/neutrophil, p<0.001; 65.1 ± 5.85 % internalization of total neutrophil associated NPs, p<0.001) compared to uncoated NPs (0.76 ± 0.25 internalized NPs/cell; 10.95 ± 3.22 % internalization of total neutrophil associated NPs, Figure 1C-G).

### BSA-NP association with neutrophils did not inhibit their activation *in vitro*

Our prior studies have utilized nitric oxide (NO) level as an indicator of neutrophil activation following cabozantinib treatment, which correlates with neutrophil mediated anti-tumor immune responses in Pb-Cre;PTEN^fl/fl^p53^fl/fl^ mice (8). To test the hypothesis that nanoparticle internalization within neutrophils does not negatively impact their activation, we incubated BSA-NPs with neutrophils in conditioned media harvested from cabozantinib-treated murine PTEN/p53-deficient tumor-derived SC1 cells. We observed significantly increased NO levels following incubation with conditioned media (225.7±13.44 at 60 minutes relative to 89.38±3.9 at baseline, p<0.05) in unloaded murine neutrophils within 60 minutes, which was sustained up to 3 hours. When neutrophils were incubated with a 100 μg dose of BSA-NP containing conditioned media, DiR-loaded BSA-NP association (p<0.05) reached a plateau within 120 minutes, following neutrophil activation. NO levels were not changed in neutrophils before (225.7±13.44) and after (182.4±29.93) association of DiR-loaded BSA-NP (Figure 2A and C). Additionally, NO staining was similar among DiR-loaded BSA-NP associated neutrophils with different quantities of associated BSA-NPs (Figure 2A and B). Taken together, these data indicate that BSA-NP association and internalization by neutrophils, does not alter their activation status.

**Figure 2.**
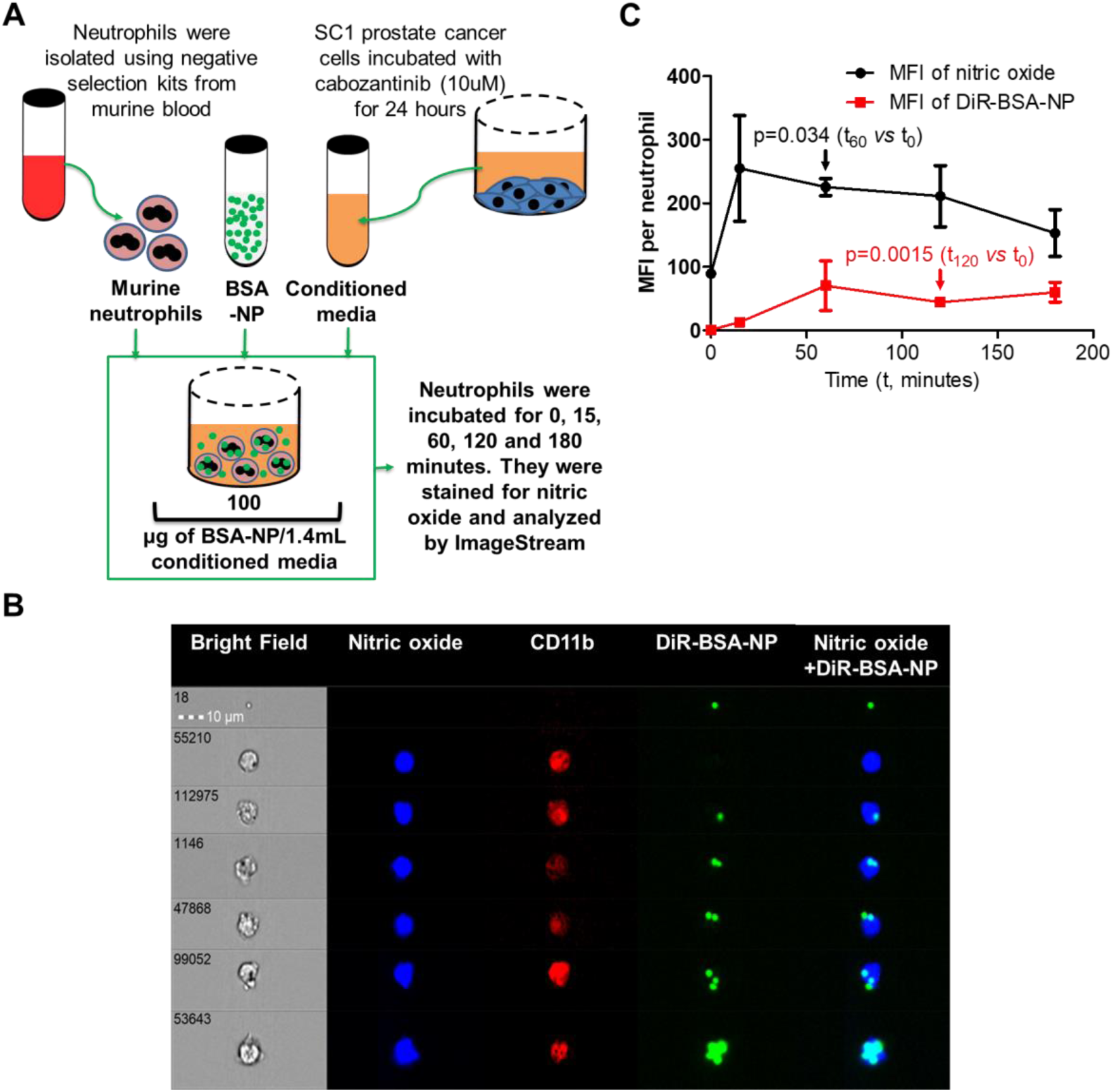
BSA-NP binding with murine neutrophils does not alter their activation status. (A) Experimental schema showing BSA-NP-associated neutrophil activation *in vitro* by nitric oxide (NO) staining. Murine SC1 prostate cancer cells were treated with cabozantinib, and conditioned media was collected to activate neutrophils. BSA-NP were added to conditioned media and incubated with neutrophils. NO staining was performed and fluorescence intensity was determined using ImageStream in BSA-NP associated or unbound neutrophils. (B) ImageStream images showed single neutrophil event in Brightfield camera and its staining with anti-CD11b antibody by red color. NO level and associated BSA-NP per neutrophil were represented by blue and green fluorescence, respectively. Each row represented a single neutrophil, and depicts NO levels and NP uptake (C) Neutrophil activation after BSA-NP association was evaluated by determination of an average activation status (MFI of NO) and bound BSA-NP (MFI of DiR-BSA-NP) per neutrophil at different time points (0, 15, 60, 120 and 180 minutes) after incubating neutrophils with a 100μg dose of BSA-NP containing conditioned media. t indicated time in minutes; t_0_= 0 minute, t_60_= 60 minutes, t_120_= 120 minutes after incubation of BSA-NP with neutrophils in conditioned media. n=3 independent experiments.

### BSA-NP association did not alter activation or functionality across a range of doses, and NP uptake was completely saturated at 100 μg equivalent dose *in vivo*

We next tested the impact of BSA-NP doses on binding capacity and functionality of murine neutrophils. When neutrophils were incubated with 10, 100 and 1000 μg doses of DiR-loaded BSA-NP in conditioned media, a significant (p<0.05) increase in BSA-NP association with neutrophils was observed at 100 μg dose, which was sustained at 1000 μg dose of DiR-loaded BSA-NP (50.93±6.298 and 46.12±9.659, respectively), compared to the 10μg dose of DiR-loaded BSA-NP treated group (25.6±2.913, Figure 3A). None of the DiR-loaded BSA-NP doses altered nitric oxide status of activated neutrophils (Figure 3B). Furthermore, cytokine array analysis of conditioned media after incubation of neutrophils with 0, 10, 100 and 1000μg doses of BSA-NP, demonstrated a selective increase in neutrophil-secreted cytokines, CCL2 and CXCL10, following activation. DiR-loaded BSA-NP did not alter secretion of these cytokines from neutrophils (Figure 3C and 3D, respectively) at any NP doses tested, suggesting no alteration in neutrophil functionality with NP binding/internalization. Consistent with these observations, neutrophil phagocytic capacity was also not affected by any doses of DiR-loaded BSA-NP (Figure 3E). Also, there was no statistical difference in MFI of DiR-loaded BSA-NP per neutrophil between the 100 and 1000 μg doses of BSA-NP, indicating that neutrophils were bound and completely saturated with BSA-NP at the 100μg dose *in vitro*. Furthermore, the BSA-NP doses were re-suspended in 1.4 mL (equivalent to average blood volume per mouse) of conditioned media in these *in vitro* experiments to mimic NP concentrations achieved *in vivo*. These data demonstrate that 100μg of BSA-NP is the optimal dose to achieve maximum association of BSA-NP with neutrophils *in vivo*.

**Figure 3.**
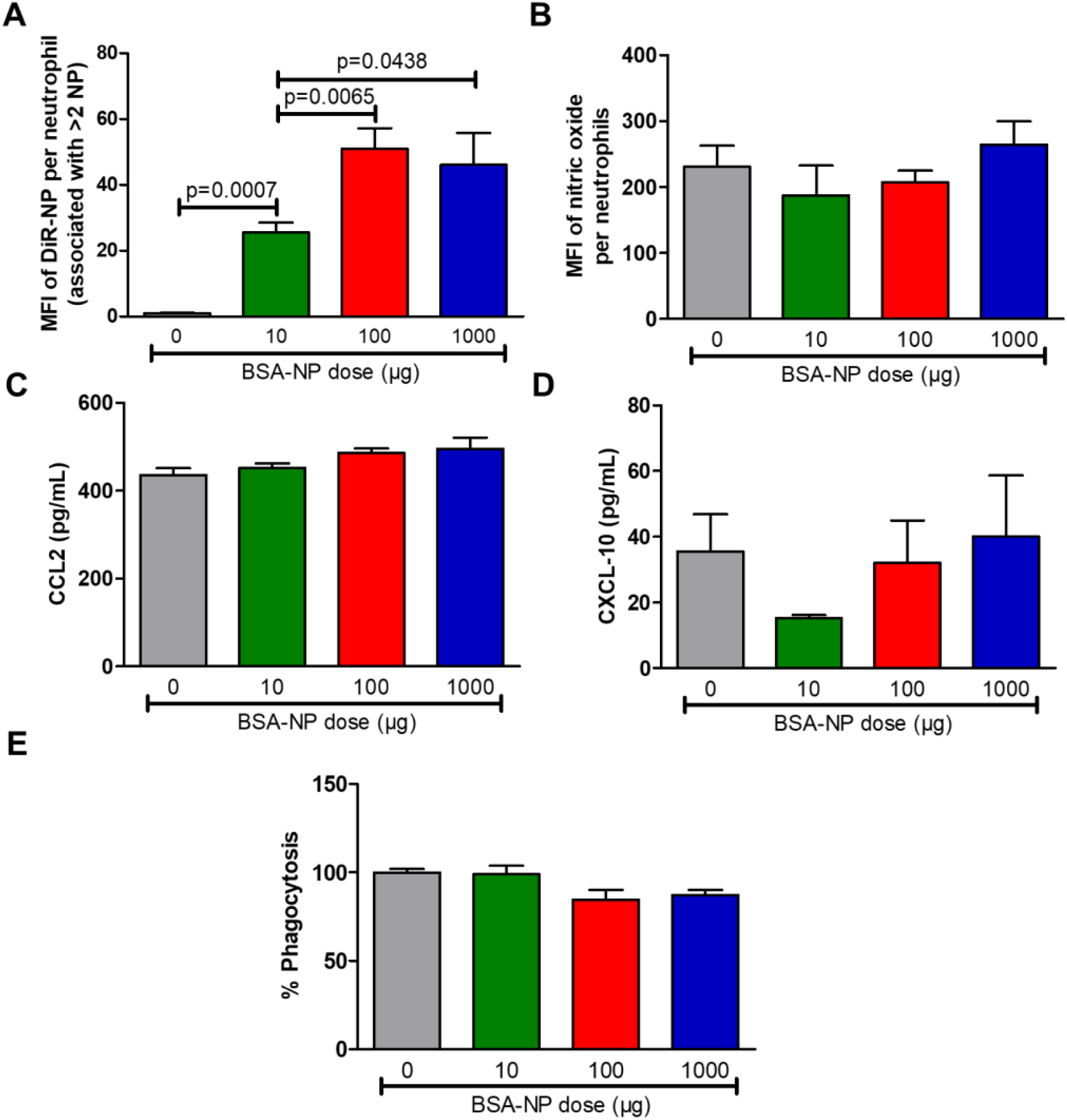
Maximum association of BSA-NP with neutrophils achieved at 100 μg *in vivo* equivalent dose, without alteration in neutrophil activation/functionality *in vitro*. Murine neutrophils were incubated with 0, 10, 100 or 1000μg of BSA-NP containing SC1-conditioned media for 3 hours and their BSA-NP binding capacity, (MFI of DiR-BSA-NP per neutrophil, A), and activation status (MFI of NO per neutrophil, B), were assessed. Furthermore, 22 cytokines were estimated using LEGENDplex™ bead-based immunoassay in conditioned media after 6 hours incubation of neutrophils with 0, 10, 100 and 1000μg doses of BSA-NP. Neutrophil secreted cytokines, CCL2 (B) and CXCL10 (D) are shown. For phagocytosis assay (E), SC1 cells were treated with cabozantinib (10uM) for 24 hours and stained with DiI dye, and then incubated with neutrophils for 6 hours in presence of 0, 10, 100 and 1000μg of BSA-NP. To calculate percent phagocytosis, the uptake of DiI dye per neutrophil was monitored using flow cytometry. n=3 independent experiments.

### Cabozantinib enhanced *in vivo* delivery of BSA-NPs to prostate tumors via a neutrophil-specific mechanism

We have previously shown that cabozantinib activates neutrophil-mediated anti-cancer innate immunity within 72 hours in Pb-Cre; PTEN^fl/fl^, p53^fl/fl^ mice (8,21). Therefore, we tested the impact of cabozantinib on intratumoral delivery of dye-loaded BSA-NPs that can be internalized by neutrophils. Mice were either untreated or pre-treated with cabozantinib for 1 hour prior to the intravenous injection of dye-loaded uncoated NPs or BSA-NPs for three days. Activated neutrophils have a short half-life (6-8 hours), which could represent a barrier for optimal intratumoral delivery of NPs. To circumvent this issue, we administered twice daily dosing of NPs for 72 h and then assessed biodistribution of injected NPs via *ex vivo* fluorescence imaging of organs harvested from mice. We observed that prostate tumors from mice that were pre-treated with cabozantinib followed by BSA-NP exhibited high fluorescence uptake, relative to other organs within the same mouse and prostate tumors from untreated control mice or mice treated with uncoated NP and cabozantinib (Figure 4A-C). To determine whether the increased dye-loaded, BSA-NP delivery was mediated via a neutrophil-specific mechanism, mice were pre-treated with both cabozantinib and Ly6G antibody, which systemically depletes neutrophils. Concomitant pre-treatment with cabozantinib and Ly6G antibody abrogated the increased dye-loaded BSA-NP delivery observed with cabozantinib pre-treatment alone (Figure 4D). Quantitative analysis performed using ImageJ revealed a significant increase in mean fluorescence uptake between tumors of mice treated with cabozantinib/BSA-coated NPs (53.67+/-5.4/px^2^) vs. untreated mice (1.73+/-1.3/px^2^) or mice treated with either cabozantinib/uncoated NP (14.47+/-3.8/px^2^) or cabozantinib/BSA-NP/Ly6G antibody (4.93+/-2.5/px^2^) (p < 0.05) (Figure 4E). These results demonstrate that cabozantinib enhanced *in vivo* delivery of BSA-NP to PTEN/p53-deficient murine prostate tumors via a neutrophil-specific mechanism.

**Figure 4.**
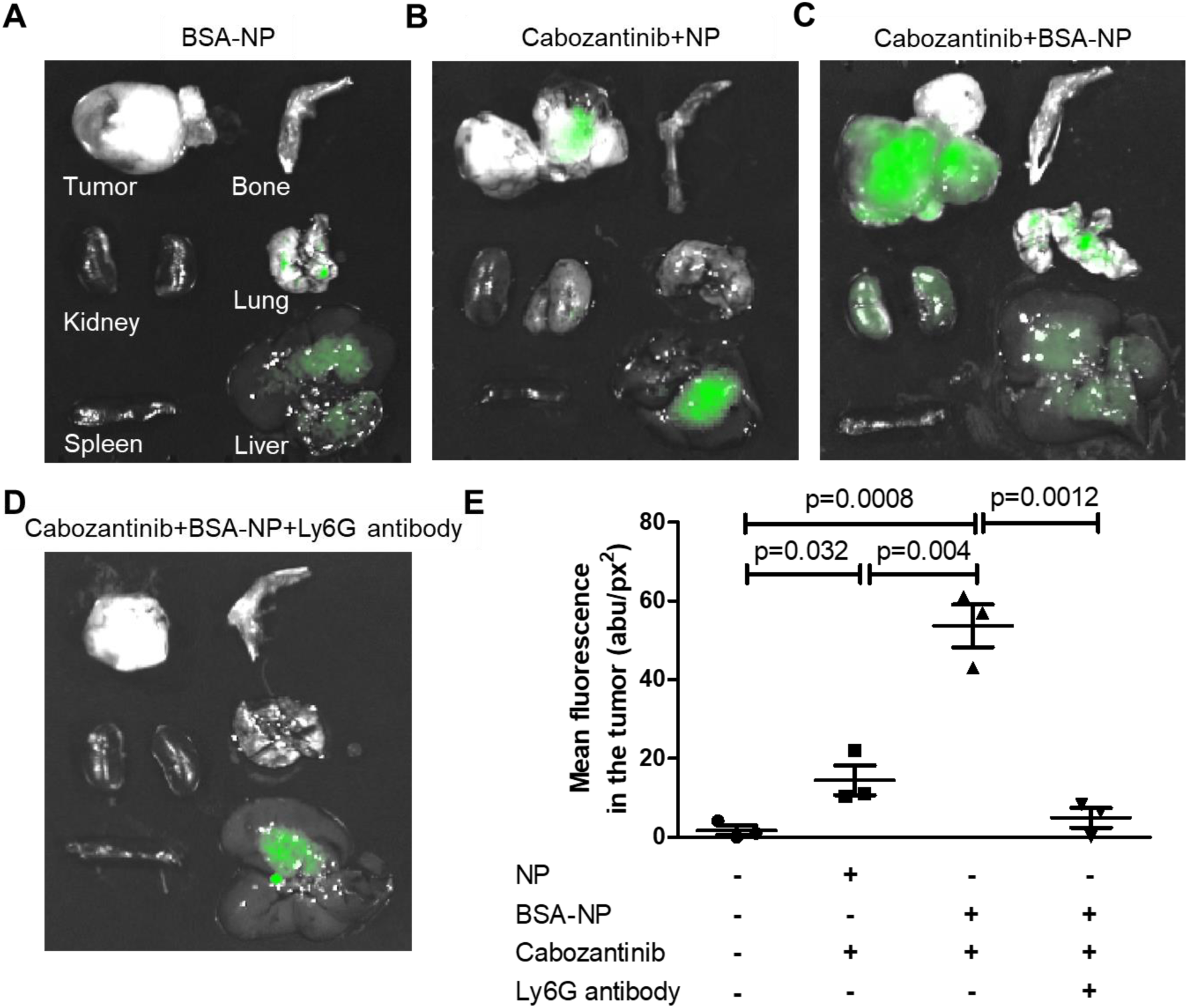
Cabozantinib treatment *in vivo* results in an increase in neutrophil-associated, dye-loaded BSA-NPs within prostate tumors. (A) Pb-Cre; PTEN^fl^p53^fl/fl^ mice (n=3 mice per group) with established prostate tumors, were treated with BSA-NP alone, cabozantinib plus uncoated NP, cabozantinib plus BSA-NP cabozantinib plus BSA-NP plus Ly6G antibody, as described in Methods Tumor, kidney, spleen, bone, lung and liver were harvested from each mouse and IVIS imaging were performed to assess NP uptake in these organs. The fluorescence images of these organs were normalized using background fluorescence from respective non-NP treated organs on the IVIS system. The overlay of fluorescent images with bright field images were done on IVIS system and presented for (A) BSA-NP (B) cabozantinib+uncoated NP, (C) cabozantinib+ BSA-NP (D) cabozantinib+BSA-NP+Ly6G antibody (neutrophil depletion) treated groups. (E) Tumors were delineated on overlaid fluorescence/bright field images using image J, and mean fluorescence was obtained to evaluate NP uptake.

### Flow cytometry analysis revealed an increase in neutrophil-associated, dye-loaded BSA-NPs within prostate tumors of cabozantinib-treated mice

To directly quantify the association of neutrophil infiltration with enhanced delivery of DiO-loaded NP *in vivo*, tumors were analyzed for co-association of DiO-NP and neutrophil markers by flow cytometry. Consistent with our prior published work, we observed an increase in CD11b+Gr-1+ cells within the tumor, which represent tumor-infiltrating neutrophils and polymorphonuclear myeloid derived suppressor cells (PMN-MDSCs) from a baseline of 20.97% of the total cells sampled by flow cytometry in an untreated control mouse, to 65.1% (p<0.001) in a representative mouse treated with cabozantinib for 72h. The CD11b+Gr-1+ population in the tumor decreased to 6.3% (p<0.001) for a representative mouse treated with cabozantinib and Ly6G antibody, confirming the opposing effects of cabozantinib and Ly6G antibody on neutrophil infiltration within the tumor (Figure 5A). Furthermore, we assessed the presence of DiO-BSA-NPs exclusively in association with neutrophils within prostate tumors; free NPs were excluded from the analysis, as they could not be retrieved during the process of preparing the tumor for flow cytometry. Consistent with the fluorescence imaging data in Figure 4, we observed an increased frequency of CD11b+Gr-1+ cells within the prostate tumors that associated with DiO-BSA-NPs in the cabozantinib-treated mice (77.37%, p<0.001), relative to untreated mice (16.47%, Figure 5B). When mice were treated with uncoated NP and cabozantinib, the proportion of CD11b+Gr1+ cells (58.67±3.71%) were increased in the tumor, similar to BSA-NP+cabozantinib treated group (65.1±5.86%, Figure 5A). However, the frequency of uncoated NP associated CD11b+Gr1+ cells was ∼3 fold lower at 24.55% in the tumor following cabozantinib treatment, relative to BSA-NP+cabozantinib treated mice (77.37%, Figure 5B). Thus there is an increase in the proportion of CD11b+Gr-1+ cells in the tumor as well as the proportion of NP-associated neutrophils following treatment with cabozantinib only when the NPs are coated with BSA. Consistent with data shown in Fig. 4, mice treated with both Ly6G antibody and cabozantinib had the lowest frequency of DiO-associated-CD11b+Gr1+ cells (9.4%, p<0.001, Figure 5B). Taken together, cabozantinib treatment resulted in significantly increased neutrophil infiltration within PTEN/p53 deficient prostate tumors, and nearly 80% of these CD11b+Gr-1+ cells were associated with DiO-loaded BSA-NPs. The BSA-NP delivery was neutrophil-mediated, as neutrophil depletion with Ly6G pre-treatment abrogated NP delivery to the tumor in mice treated with cabozantinib. Next, we determined the % NP uptake using fluorometry, which revealed that the total uptake of BSA-NP was 0.11%, which was significantly increased by cabozantinib treatment to 0.96% (p=0.002) and decreased by Ly6G-mediated neutrophil depletion to 0.03% (p=0.0011, Figure 5C).

**Figure 5.**
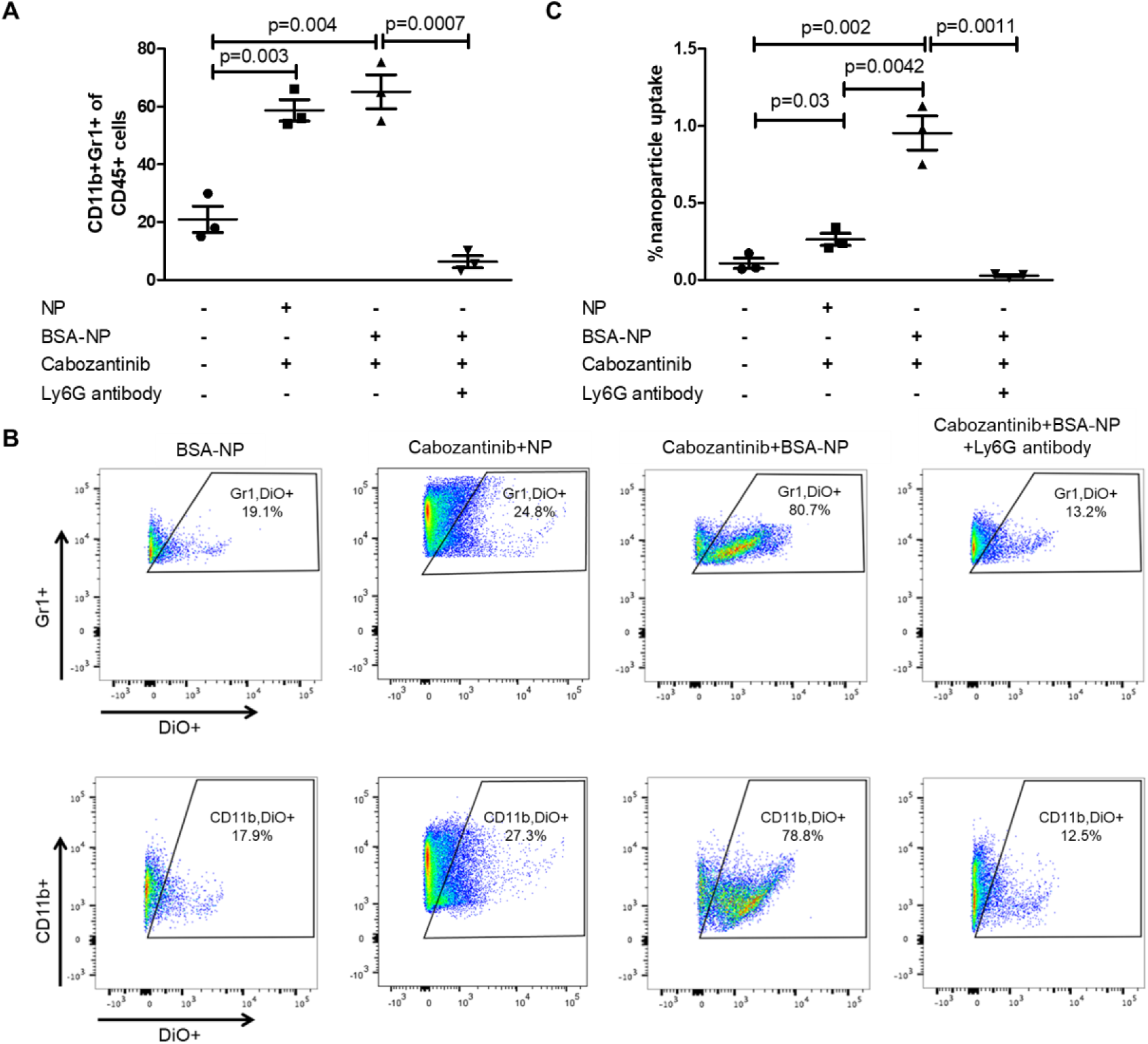
Cabozantinib increases neutrophil-associated BSA-NP uptake into the prostate tumor microenvironment. Prostate tumors of Pb-Cre; PTEN^fl^p53^fl/fl^ mice (n=3 mice per group) were treated with BSA-NP, cabozantinib+uncoated NP, cabozantinib+BSA-NP and cabozantinib+BSA-NP+Ly6G antibody for 3 days, as described in Figure 4. (A) Tumors were harvested and single cell suspensions were stained with anti-CD11b antibody, anti-Gr1 antibody and frequency of tumor-infiltrating neutrophil were analyzed using flow cytometry. The CD11b+Gr1+ populations were further gated for DiO-NP. (B) Representative flow plots showing frequency of DiO-NP-associated neutrophils within the TME. (C) To determine % NP uptake within the TME, the harvested prostate tumors were homogenized, and concentration of DiO-NP was determined by fluorometry analysis (λ_excitation_= 485 nm; λ_emission_=510 nm). % NP uptake was calculated on the basis of administered NP dose.

To determine whether neutrophil-mediated BSA-NP association and internalization differentially occurs within the periphery vs. tumor microenvironment (TME,) we analyzed circulating and splenic neutrophils using flow cytometry to determine the frequency of BSA-NP-associated neutrophils, following 72 hours of treatment with BSA-NP alone or cabozantinib/BSA-NP. We observed that a small fraction of circulating (0.65±0.11%) and splenic (0.21±0.01%) neutrophils associated with BSA-NP in untreated mice, which was slightly enhanced to 1.46±0.08% (p<0.01) and 0.37±0.03% (p<0.05), respectively, in mice treated with cabozantinib (Figure 6). However, this was a very small fraction of the 80% BSA-NP-associated neutrophils harvested from the TME following cabozantinib treatment. Collectively, these data demonstrate that BSA-NPs are selectively internalized by tumor-infiltrating neutrophils (TINs) following cabozantinib treatment. A schematic of our proposed mechanism is shown in Figure 6C.

**Figure 6.**
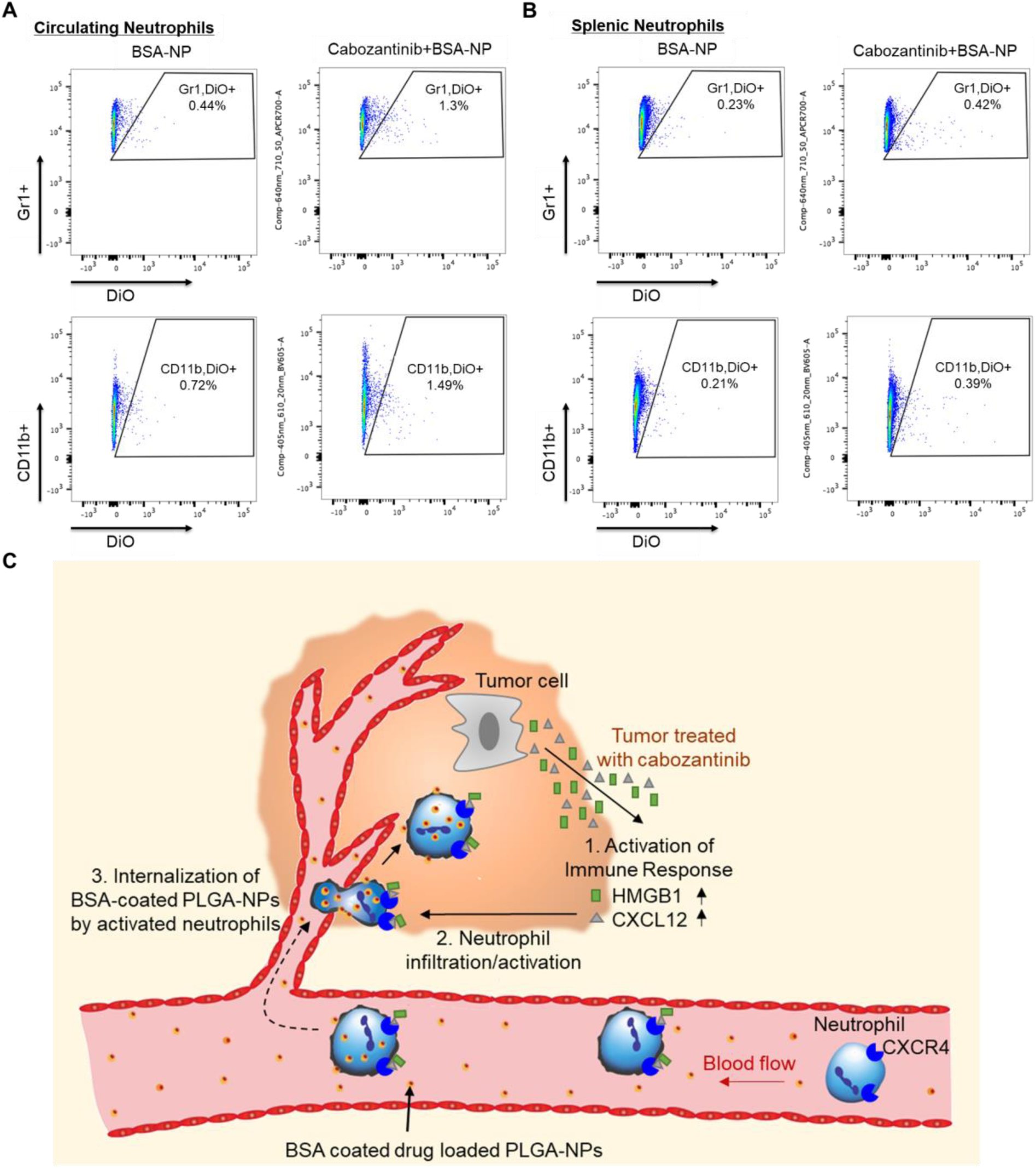
BSA-NPs do not significantly associate with blood or splenic neutrophils in response to cabozantinib treatment *in vivo*. Pb-Cre; PTEN^fl^p53^fl/fl^ mice (n=3 mice per group) with established prostate tumors, were treated with BSA-NP and cabozantinib+BSA-NP for 3 days, as described in Figure 4. Single cell suspensions were prepared from peripheral blood and splenic tissue, and stained with anti-CD11b antibody and anti-Gr1 antibody. The frequency of NP-associated neutrophils were quantified by gating CD11b+Gr1+DiO+ populations on flow cytometry. The representative flow plots showed frequency of DiO-NP associated circulating neutrophils (A) and splenic neutrophils (B). (C) Proposed model for cabozantinib-induced accumulation of BSA-coated PLGA NPs within prostate tumors. Cabozantinib enhances neutrophil activation, resulting in increased internalization of BSA-NPs within the prostate TME. This platform provides a novel nanoimmunotherapeutic strategy for enhanced therapeutic payload delivery within tumors.

## Discussion

In this study, we discovered that cabozantinib, a multi-receptor tyrosine kinase inhibitor, enhanced the intratumoral delivery of BSA-NPs into prostate tumors that develop in the context of prostate-specific PTEN and p53 deletion. The Pb-Cre; PTEN^fl/fl^p53^fl/fl^ mouse is an invasive and locally aggressive prostate cancer mouse model, that recapitulates features of advanced mCRPC (22). Mice treated with cabozantinib daily, and six injections of BSA-coated, dye-loaded PLGA NPs for 72 hours (two injections were administered per day at interval of 12 h), demonstrated an increase in mean fluorescence uptake within the tumor and NP delivery 0.96% relative to untreated controls (0.11%). The enhanced BSA-NP uptake with cabozantinib was reversed when neutrophils were systemically depleted with Ly6G antibody, thus demonstrating a neutrophil-dependent mechanism for BSA-NP delivery into the tumor. Furthermore, flow cytometry analysis demonstrated an increased association of NPs with tumor-associated CD11b+ Gr-1+ cell populations, suggesting that the BSA-NPs were associating with activated neutrophils within the prostate TME. In contrast, intratumoral CD11b+Gr-1+ neutrophils were significantly lower in mice treated with Ly6G antibody, resulting in a decrease of BSA-NP delivery to the tumor. This study demonstrates that the injection of BSA-NPs following treatment with cabozantinib can enhance the delivery of NPs to prostate tumors through a neutrophil-dependent mechanism. This nano-immunotherapeutic strategy achieved approximately 1% delivery of the systemically administered BSA-NPs into the prostate tumor, which is significantly higher than the 0.7% (median) of tumor targeted BSA-NP delivery reported in the literature to-date (11,16).

We have demonstrated that approximately 80% of CD11b+/Gr-1+ cells within the TME are associated with BSA-NPs. There are several explanations to account for the 20% of CD11b+/GR-1+ cells that are not associated with BSA-NPs. First, the PTEN/p53-deficient prostate tumors have a high frequency of myeloid-derived suppressor cells within the TME, which are indistinguishable from neutrophils by flow cytometry. Second, pharmacokinetic factors that include a 1-hour lag time between the initial administration of cabozantinib and subsequent administration of the BSA-NPs, can be improved in future studies. Third, at steady state, a pool of neutrophils are likely already present within the TME prior to administration of cabozantinib. Despite these limitations, we observed a 4-fold increase in neutrophils associated with BSA-NPs within cabozantinib-treated prostate tumors relative to untreated tumors, which represents a significant enhancement of intratumoral NP delivery. Given our previous study demonstrating that cabozantinib enhances neutrophil infiltration within PTEN/p53-deficient prostate GEMM tumors, combined with other work demonstrating internalization of NPs by activated neutrophils, we propose that the BSA-coated polymeric NPs are associating with and internalized by TINs, resulting in enhanced retention and delivery of NPs within prostate tumors (8,15). While we have not excluded the possibility that neutrophils activated within the TME following cabozantinib treatment, are recycled back into the periphery where they may associate with and internalize NPs, the data in Fig. 6 suggests that NP uptake and internalization predominantly occurs within TINs.

In this study, we have demonstrated that BSA-coating of PLGA NPs promotes association with and internalization by activated neutrophils. This is consistent with prior studies demonstrating that NPs made from denatured BSA could target activated neutrophils *in situ* and deliver therapeutics across blood vessel walls (15). Intravenously injected BSA NPs were preferentially internalized by activated neutrophils and were able to cross the blood vessel wall in response to inflammation (15). A follow-up study evaluated an application of this principle and successfully demonstrated that drug-loaded, BSA-NPs, when co-administered with TA99, a monoclonal antibody specific for gp75 antigen, resulted in enhanced neutrophil activation, recruitment and tumor growth inhibition via an antibody-dependent cellular cytotoxicity mechanism, relative to NPs or TA99 alone (16). These experiments reflect the potential of harnessing neutrophil activation to promote efficient NP delivery to tumors. In our study, we integrate a novel tyrosine kinase inhibitor mediated neutrophil activation strategy to enhance BSA-coated PLGA-NPs uptake within the prostate tumors that develop in an aggressive, treatment-refractory mouse model of advanced PCa.

In our previous study, we demonstrated that cabozantinib-mediated neutrophil infiltration lead to tumor clearance in prostate-specific PTEN/p53 GEMMs (8). Our findings in this study offer a potential anti-cancer strategy that integrates nanomedicine and cancer immunotherapy via activation of innate immunity. We have demonstrated that neutrophil internalization of BSA-coated drug-loaded PLGA NPs has no effect on neutrophil activation, which is consistent with prior studies that have demonstrated that BSA-NP uptake does not affect the mobility, activation or cytokine release of neutrophils (15), and that PLGA NPs are not cytotoxic to neutrophils (23). This study opens the door to further investigation into next-generation NP-targeting technologies. We used larger BSA-NPs (450 nm), relative to ∼180 nm diameter BSA-NPs used in previous tumors targeted delivery studies (15,16,20), as the 450 nm particles can achieve higher drug loading and the reduced surface area to volume ratio enables longer controlled release. Future studies are needed to evaluate drug release kinetics of large (450 nm) vs small (100-200 nm) BSA-NPs.

To maximize the potential clinical translation of this technology, it may also be possible to optimize this targeted approach by altering the BSA-coating – for example, by using denatured BSA rather than native BSA used in this study, and screening for additional tyrosine kinase inhibitors or precision medicine therapies which promote neutrophil activation and infiltration. Greater neutrophil infiltration has been associated with improved survival in early stage lung cancer and gastric cancer (24-26), highlighting the broad applicability of a neutrophil-mediated targeted NP delivery strategy in oncology. Furthermore, PCa most commonly metastasizes to the bone, and recent studies have shown that neutrophils are enriched within tumor areas in bone metastatic PCa patients. As metastatic prostate tumors evolve within the bone tumor microenvironment, the cancer cells evade neutrophil-mediated cell killing (13,27). Taken together, these data underscore the potential of cabozantinib-mediated neutrophil activation and intratumoral drug-loaded NP delivery, to have specific anti-cancer activity within bone metastases.

From a translational standpoint, this activated neutrophil-based platform technology can be deployed to test novel combinatorial therapeutics to enhance intratumoral payload delivery. Given our findings that cabozantinib-induced neutrophil activation/NP uptake occurs within TINs and not peripheral neutrophils, this platform would allow for selective delivery of therapeutic payload to tumors, while mitigating non-specific organ toxicity encountered with systemic targeted and chemotherapies. This is particularly relevant in the era of combinatorial targeted therapies, such as combination of kinase inhibitors and/or chemotherapy, which can result in profound systemic toxicity in advanced cancer patients. In summary, this convergent approach has the potential to harness drug-induced innate immunity and neutrophil-mediated nanomedicine delivery for effective and safe anti-cancer therapy.

## Acknowledgements

We are thankful to Dr. Rui Kuai for providing technical comments on experiments and manuscript.

